# Development of Visually Indistinguishable Acoustic Coupling Pads for Double-Blind Focused Ultrasound Neuromodulation Studies

**DOI:** 10.64898/2026.01.27.702113

**Authors:** Samantha F. Schafer, Norman M. Spivak, Andrew A.E.D. Bishay, Alexander Bystritsky, Peter A. Lewin, Mark E. Schafer

## Abstract

**Background:** Transcranial focused ultrasound (tFUS) is an emerging noninvasive neuromodulation modality with the ability to target deep brain structures with high spatial precision. Despite its promise, rigorous evaluation of its efficacy is limited by the absence of reliable, fully double-blind sham methodologies.

**Objective:** To develop and validate a pair of visually and mechanically indistinguishable acoustic coupling pads that enable true double-blind tFUS neuromodulation studies by providing either efficient ultrasound transmission or robust ultrasound blocking without altering participant or operator experience.

**Methods:** Two coupling pads were engineered: a transmitting pad designed to allow <5% pressure amplitude loss relative to free-water propagation, and a non-transmitting pad designed to attenuate ultrasound by ≥40 dB. Both pads used a Dragon Skin 10 NV silicone base and were identical in size, appearance, flexibility, and handling. The non-transmitting pad incorporated an encapsulated air-based blocking layer using an open-cell polyethylene foam insert. Acoustic performance was evaluated in a water tank using a 650 kHz BrainSonix transducer and a calibrated needle hydrophone. Sound speed of the silicone material was measured using pulse-echo techniques.

**Results:** Twenty-three matched transmitting and non-transmitting pad pairs were fabricated and tested. Transmitting pads demonstrated a mean attenuation of −0.41 ± 0.53 dB, satisfying the design criterion of minimal acoustic loss.Non-transmitting pads demonstrated a mean attenuation of −48.61 ± 4.33 dB, exceeding the required −40 dB threshold for effective sham conditions. The Dragon Skin 10 NV substrate exhibited a sound speed of 964.72 m/s and produced <2 mm axial focal shift for standard pad thicknesses, with no measurable change in focal width. Both pad types were visually and tactually indistinguishable, could not be differentiated by experienced operators or participants, and maintained mechanical integrity after repeated cleaning

**Conclusion:** These acoustically engineered coupling pads provide a practical and validated solution for achieving true single- and double-blind conditions in tFUS neuromodulation studies. By preserving identical sensory and procedural experiences while enabling precise control over ultrasound transmission, this approach addresses a critical methodological gap in human ultrasound neuromodulation research.

## Introduction

This work presents the design, fabrication, and validation of visually indistinguishable transmitting and non-transmitting coupling pads that enable reproducible, double-blind focused ultrasound neuromodulation studies. Focused ultrasound is a novel brain stimulation method with certain key advantages over other brain stimulation modalities: it’s entirely noninvasive unlike deep brain stimulation which requires surgery; it can target deep brain structures unlike transcranial magnetic stimulation (TMS) and transcranial direct current stimulation (tDCS) which are surface-limited; it can inhibit or excite neurons depending on ultrasound exposure settings; and it offers better spatial resolution (< 1cm) compared to TMS or tDCS, which affect larger regions.

All medical ultrasound applications, including focused ultrasound for neuromodulation, require a completely air-free mechanical coupling between the transducer contact surface and the participant’s skin. This is because there is a substantial acoustic impedance (defined as the product of medium density and speed of sound) mismatch between air and the transducer or skin, making air highly reflective (R= 99.9%) [1, 2]. If a practitioner placed a diagnostic imaging probe directly on a participant without any coupling medium, even a thin air layer between the probe and skin would be sufficient to block ultrasound propagation (or transmission), resulting in no image being displayed on the imaging device. Therefore, all ultrasound studies require the use of a coupling medium; the most common is a viscous, water-based gel, although alternatives such as acoustic standoff pads are also used.

Any well-designed clinical trial involves assessing treatment results against a placebo or control group to determine statistical significance, most commonly through blinding (ideally double blinding) [3]. However, bias, whether intentional or unintentional, can still occur depending upon the method of blinding used, and can affect the significance of the study results [3]. Proper blinding throughout the study also helps minimize the placebo effect [3]. Studies utilizing ultrasound for neuromodulation are no exception and require a sham or blinding method to reinforce results and assess the efficacy of this emerging technology.

During ultrasound exposure, the transducer can generate signals in the audible frequency range, typically at the pulse repetition frequency. This is discernable to the study subject as a clicking or buzzing sound. In addition, there may be vibrations that are coupled to the subjects’ bone and thereby conducted to the auditory or vestibular nervous systems. Thus there are several components of the ultrasound treatment experience which must be considered when looking to create non-exposure or “sham” conditions.

Creating sham conditions and adequate blinding of focused ultrasound is thus of key importance to evaluating its potential as a neuromodulatory technique. Unfortunately, no good sham conditions exist yet. Several methods have been trialed, although none of them are ideal. The first method involves a simple “on/off” protocol where the active group receives treatment with an energized transducer, while the sham group does not because the transducer remains unenergized. This approach would not produce the same auditory cues for participants; additionally, a designated individual must be involved to decide whether or not to energize the transducer. In the Sanguinetti et al. study, blinding was achieved via a pre-programmed interface[4]. In the Badran et al. study, a separate person operated the ultrasound system to turn it on or off [5].

The second method disrupts ultrasound access to the brain by disconnecting the transducer from the skull. Liu et al. and Yu et al. detached the transducer from participants’ skulls, removing any acoustic connection [6, 7].Legon et al. also disrupted this connection by turning the transducer over, so the non-transmitting side faced the skull instead of the transmitting side[8].

Some researchers conceal auditory cues generated by the transducer during treatments by playing engineered white noise or adaptive sound through headphones to hide confounds at the ultrasound’s pulse repetition frequency[9]. This method is often combined with a sham method to prevent placebo effects.

Sham conditions can be also created by blocking ultrasound transmission, as described by Legon et al., who used a “high acoustic impedance disk on the face of the transducer.”[10]. However, the authors did not specify the disk material or its impedance level. Although the disk was mechanically coupled to the skull, it prevented ultrasound from reaching the participant. Furthermore, the authors did not compare the physical properties of the disk to a typical coupling material like Aquagel. These details suggest the study was likely single-blind, meaning the raters were privy to the group assignment of the subjects, again reiterating the need for validated and effective double-blinding methods for tFUS.

Although existing sham methods can function effectively in theory, they depend on a carefully managed flow of information among researchers and participants, creating opportunities for bias. Because at least one staff member must know each participant’s treatment condition, subtle verbal or nonverbal cues could unintentionally influence outcomes. Participants, too, may infer their group assignment—especially if auditory or tactile sensations differ between active and sham conditions, such as when the transducer is off, flipped, or decoupled from the skull. These risks are amplified in crossover designs where subjects experience both active and sham conditions. Therefore, maintaining identical procedures, environmental cues, and participant experiences across all sessions and operators is essential to preserve study validity.

To address these limitations, a coupling approach is necessary that not only maintains identical sensory and procedural conditions between sham and active sessions but also ensures precise, controllable transmission of ultrasound energy—thus upholding both experimental rigor and true blinding integrity. This requirement drives the development of specialized acoustic coupling pads designed to standardize the interface between the transducer and scalp, allowing consistent exposure control while eliminating sensory or procedural differences between treatment conditions. This work presents the design, fabrication, and validation of visually indistinguishable transmitting and non-transmitting coupling pads that enable reproducible, double-blind focused ultrasound neuromodulation studies.

## Methods

### Pad Design Objectives

Two visually and mechanically indistinguishable acoustic coupling pads were developed for transcranial focused ultrasound (tFUS) research: a transmitting pad engineered to allow <5% pressure amplitude loss relative to free-water propagation and a non-transmitting pad designed to attenuate ≥40 dB of ultrasound energy (≥100× pressure amplitude reduction). These performance criteria ensured that the transmitting pad would function equivalently to standard coupling media while the non-transmitting pad would block ultrasound sufficiently to serve as a sham condition. This was done to enable fully double-blind neuromodulation experiments, such that neither operators nor participants could determine whether a treatment was active or sham based on tactile, auditory, or visual cues. The pads were required to be visually identical, mechanically stable, flexible, skin-safe, reusable, and capable of being sanitized with isopropyl alcohol between participants without mechanical or chemical degradation. The material needed to conform to the face of the transducer and adequately fill the gap between the transducer and the participant. Because the pads are in direct contact with the skin, they were required not to cause irritation. To satisfy these requirements, a sandwich pad configuration was developed consisting of an acoustically conductive silicone base layer with or without a central acoustically blocking layer (Figure 1). A rigid plastic alignment ring was embedded into the pad during molding to aid in consistent positioning and attachment to the transducer face. The blocking medium was located centrally within the silicone to ensure the non-transmitting pad was mechanically and visually indistinguishable from the transmitting pad.

**Figure 1:**
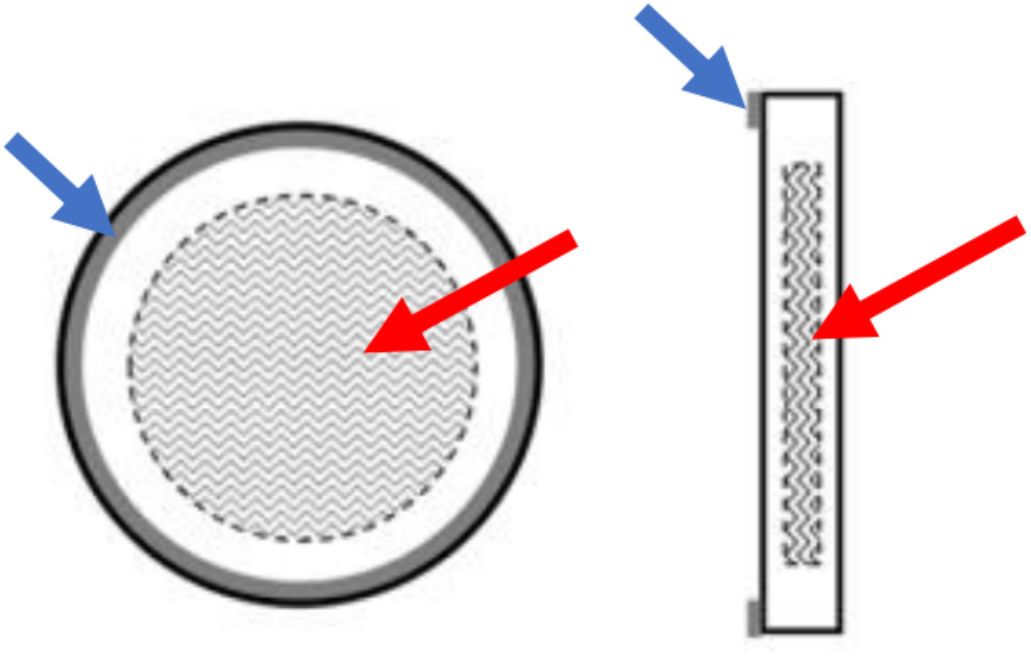
Schematic of Pad Design. Red arrow shows the central acoustically blocking layer. Blue arrows correspond to the silicone base layers sandwiching the blocking layer. The final pad is 62 mm in diameter and 5 mm thick.

### Material Screening and Selection

Candidate materials were evaluated in multiple phases. First, potential pad base materials were screened for acoustic and mechanical suitability, with acoustic attenuation given highest priority. Materials were also evaluated for manufacturability, handling characteristics, and availability from reliable commercial suppliers. Smooth-On, Inc. was selected due to its broad selection of skin-safe polymers and ready availability of material safety data sheets. Nine skin-safe silicone materials were obtained and tested (Table 1, Figure 2). Each material was mixed according to the manufacturer’s instructions and degassed in a benchtop vacuum chamber for two five-minute intervals at –90 kPa, the maximum achievable for a standard low-vacuum system. The first degassing was performed in the mixing vessel, and the second was performed after pouring into molds.

**Table 1:**
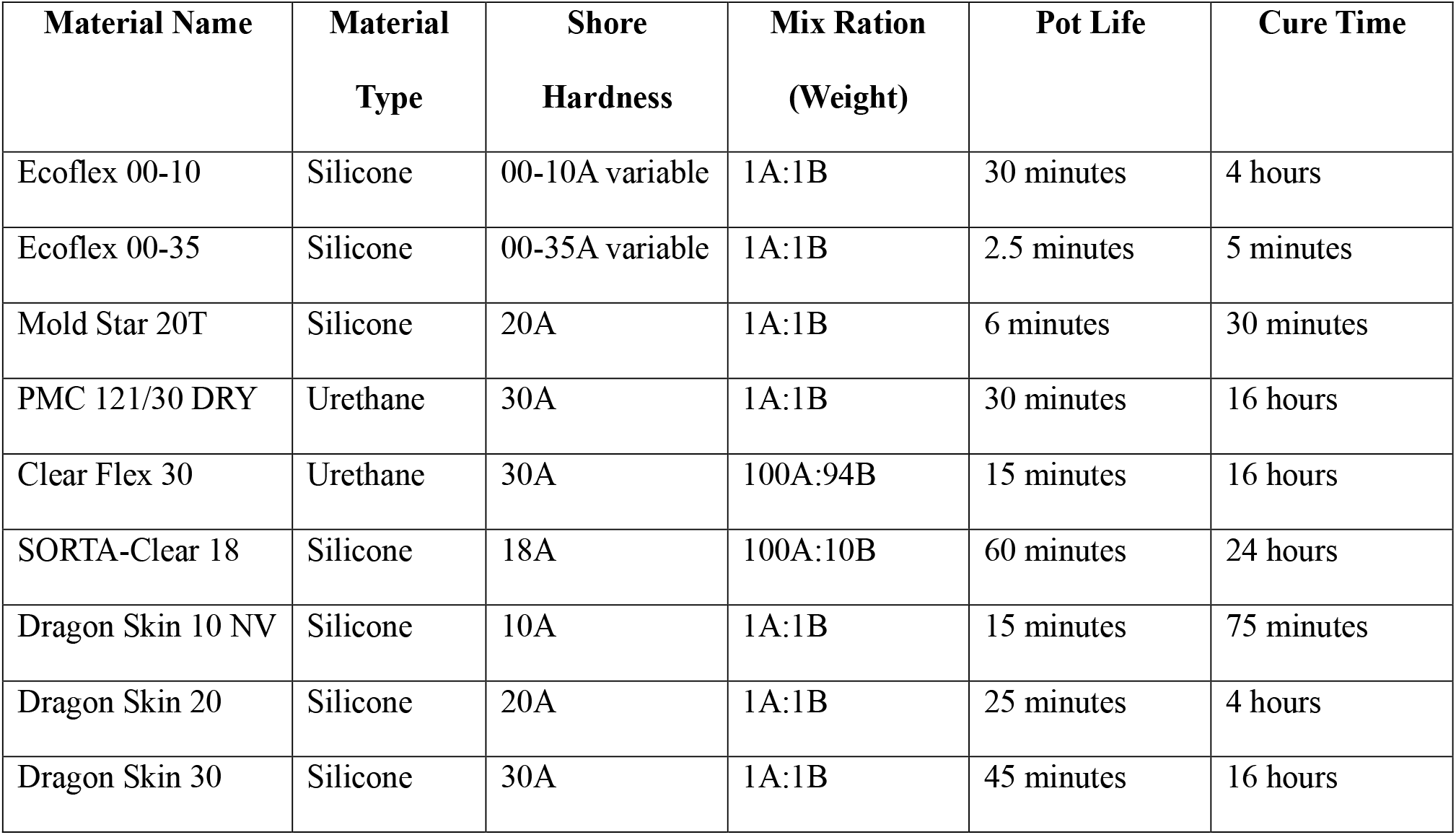
List of Materials for Acoustic Compatibility Testing and their Properties. All material characteristics correspond to an ambient room temperature of 20 ºC.

**Figure 2:**
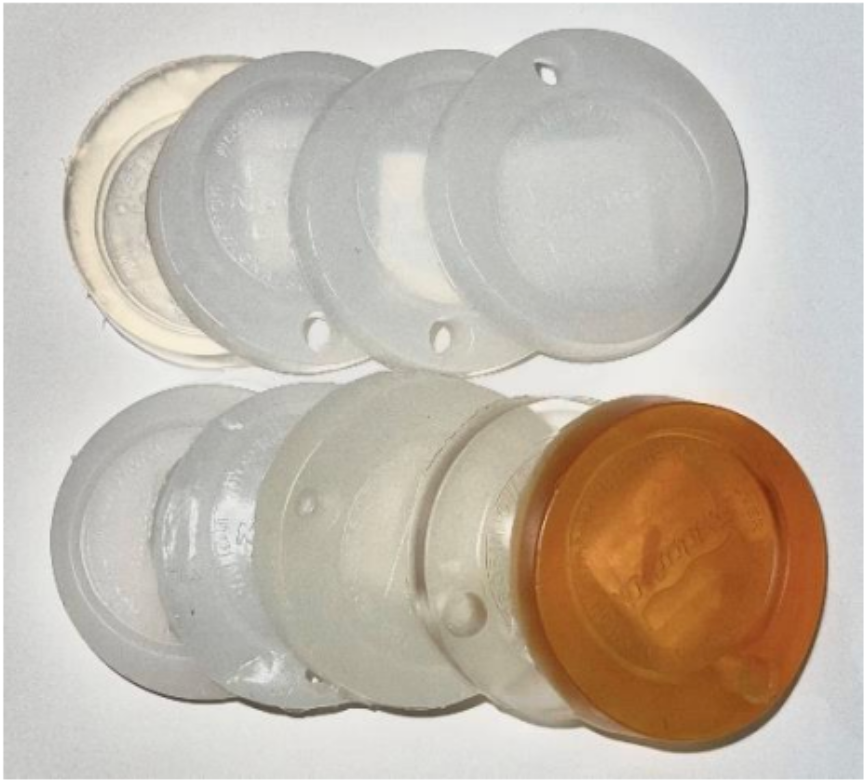
Image of coupon material samples that were used for acoustic compatibility testing

Circular coupons 53 mm in diameter and approximately 5 mm thick were cast for each material and tested in a deionized-water tank with a first generation BrainSonix transducer. The circular single element transducer was 60 mm in diameter, operating at 650 kHz, which is within the typical 500-750 kHz that is used for neuromodulation [11]. It had a nominal focal distance of 80mm, and the coupon was placed at 60mm from the transducer face to ensure that the full acoustic beam cross-section passed through the material prior to measurement. A Reson TC4038 needle hydrophone was positioned at the nominal focal distance (2cm beyond of coupon), to measure the pressure amplitude at the focal zone. With a fixed transmit voltage, peak-to-peak voltage at the output of the Reson hydrophone was recorded using a Tektronix TDS3012C oscilloscope.

Dragon Skin 10 NV silicone demonstrated the lowest attenuation (−1.04 dB compared to water), along with favorable characteristics including low Shore hardness for conformal coupling, ability to be pigmented for opacity, compatibility with repeated alcohol cleaning, and a rapid ∼75-minute cure time enabling same-day pad fabrication. Thus, Dragon Skin 10 NV was selected as the base material for all pad prototypes.

### Non-Transmitting Layer Selection and Integration

The blocking layer was required to effectively prevent ultrasound transmission while remaining thin, flexible, and fully concealed within the pad. To achieve the required ≥40 dB attenuation for sham pads, air was chosen as the blocking medium due to its extreme acoustic impedance mismatch with the piezoelectric transducer surface (reflection coefficient ≈ 1). To implement an air layer in a stable and uniform manner, a thin open-cell polyethylene foam membrane approximately 1 mm thick was chosen. A 7.62 × 7.62 cm sample of the foam was tested in the same water-tank setup described above, and the resultant signal was demonstrated >50dB loss, confirming its suitability as a blocking medium. However, because the foam floats in uncured silicone and visible or tactile differences would compromise blinding, a two-stage encapsulation method was developed.First, a ∼2.5-mm layer of Dragon Skin 10 NV was poured into an aluminum mold and left 25 minutes topartially cure. The foam insert was then placed at the pad’s mid-plane, and a second pour was performed to reach a total thickness of ∼5 mm, fully embedding the insert without exposing edges or altering surface features. Pads were pigmented opaque white (Silc-Pig, Smooth-On, Inc.) to conceal internal architecture.

### Detailed Pad Fabrication

Dragon Skin 10 NV components were mixed in equal parts, degassed twice for five minutes each at −90 kPa, and poured into molds containing the rigid plastic ring. For non-transmitting pads, the foam insert was suspended at mid-thickness using the two-stage curing approach. After curing, pads were demolded, trimmed, inspected for uniformity, and cleaned. Pads were sanitized with isopropyl alcohol between uses.

### Sound Speed Measurement

To characterize acoustic propagation properties of the pad substrate, four samples were cast: 5 mm and 10 mm disks of Dragon Skin 10 NV and Elastack, a material with sound speed comparable to water. Testing was conducted in deionized water at 20 °C in a pulse-echo configuration using a Panametrics PR02 pulser/receiver, a 2.25 MHz unfocused ultrasonic transducer with a 0.75″ aperture, and a Tektronix TDS3012C oscilloscope. Although the target frequency for neuromodulation is 500-700kHz, a higher frequency transducer was used for this measurement to improve the time resolution of the results. A Van Keuren optically flat silica reflector (surface roughness Ra ≤ 0.2 µm) was used as a reference reflector as it allowed the ultrasound to cleanly reflect off the surface of the disk and come back to the transducer with minimal distortion. Baseline time-of-flight in water was recorded, then each disk was placed over the reflector and the change in transit time was used to compute sound speed via substitution equations. The average sound speed of both Dragon Skin 10 NV samples was calculated to be approximately 964.72 m/s with a standard deviation of ± 0.013, and the variation between measured values being 1.91% difference. The average sound speed of the Elastak samples was calculated to be approximately 1435.70 m/s with a standard deviation of ± 0.029 and the variation between measured values being 2.61% difference.

### Ultrasound Transmission Testing

Ultrasound transmission testing was performed in a deionized water tank measuring 45.72 cm x 60.96 cm x 50.8 cm (Figure 3). A 650 kHz first generation BrainSonix circular disk transducer (Figure 4) was submerged with its face below the waterline (Figure 5). The focal plane was located approximately 80 mm from the transducer face. A calibrated Reson TC4038 needle hydrophone was used to measure pressure amplitude, chosen for its known sensitivity (-230db re 1V/Pa) in the operating frequency range and small aperture(physical dimension 4mm) to minimize spatial averaging. The hydrophone was oriented co-laterally with the transmitting transducer, so that its active aperture and angular response would be the same irrespective of position in either the X or Y direction. The hydrophone was scanned laterally in the focal plane to find the maximum pressure location. Pad samples (62 mm diameter) were placed between the transducer and hydrophone (60mm from the transducer, and 20mm from the hydrophone) to ensure the entire beam passed through the pad before measurement.

**Figure 3:**
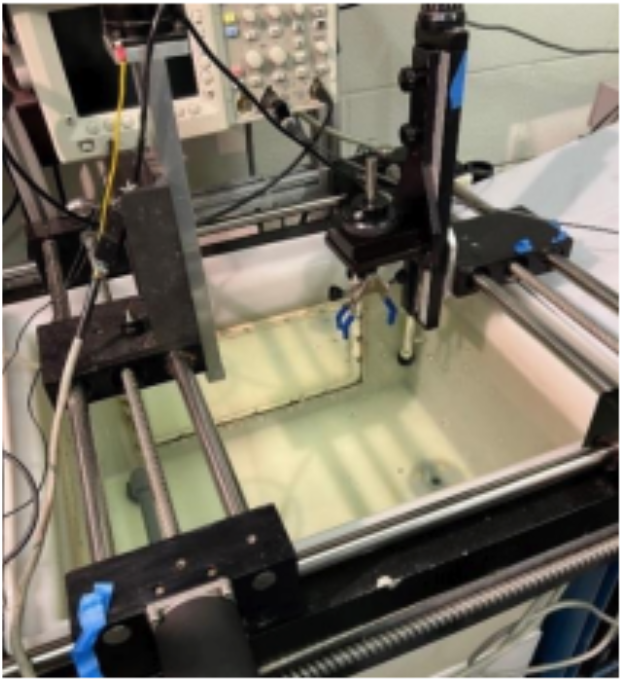
Image of water tank used for all testing.

**Figure 4:**
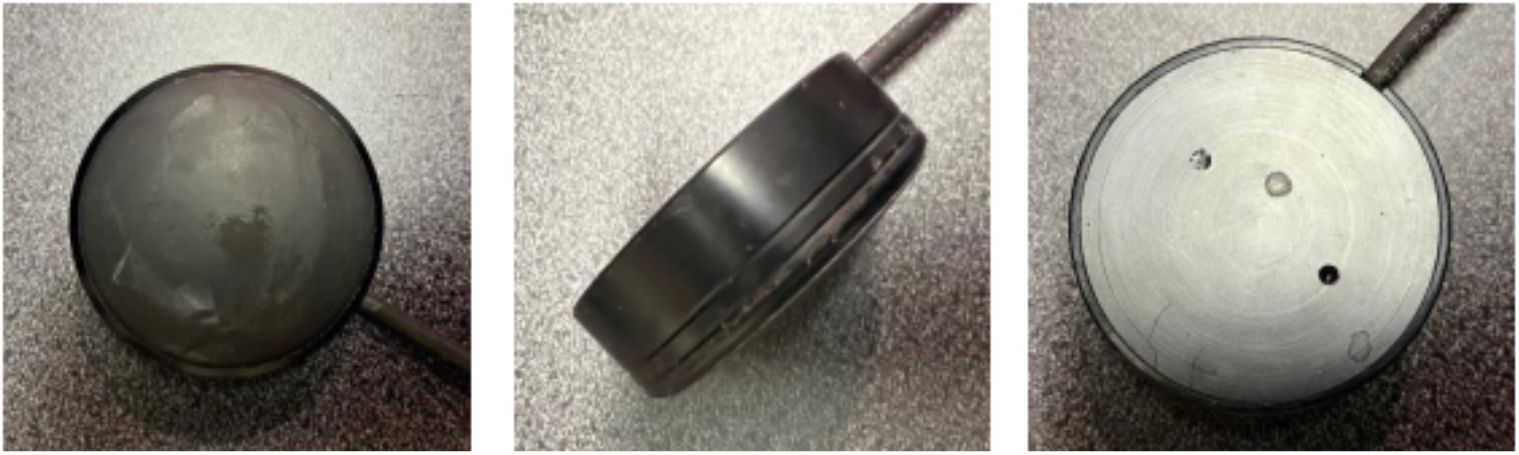
Front, side, and back views of disk transducer used for acoustic compatibility testing at 650 kHz. The transducer measures 71mm across and 27mm tall.

**Figure 5:**
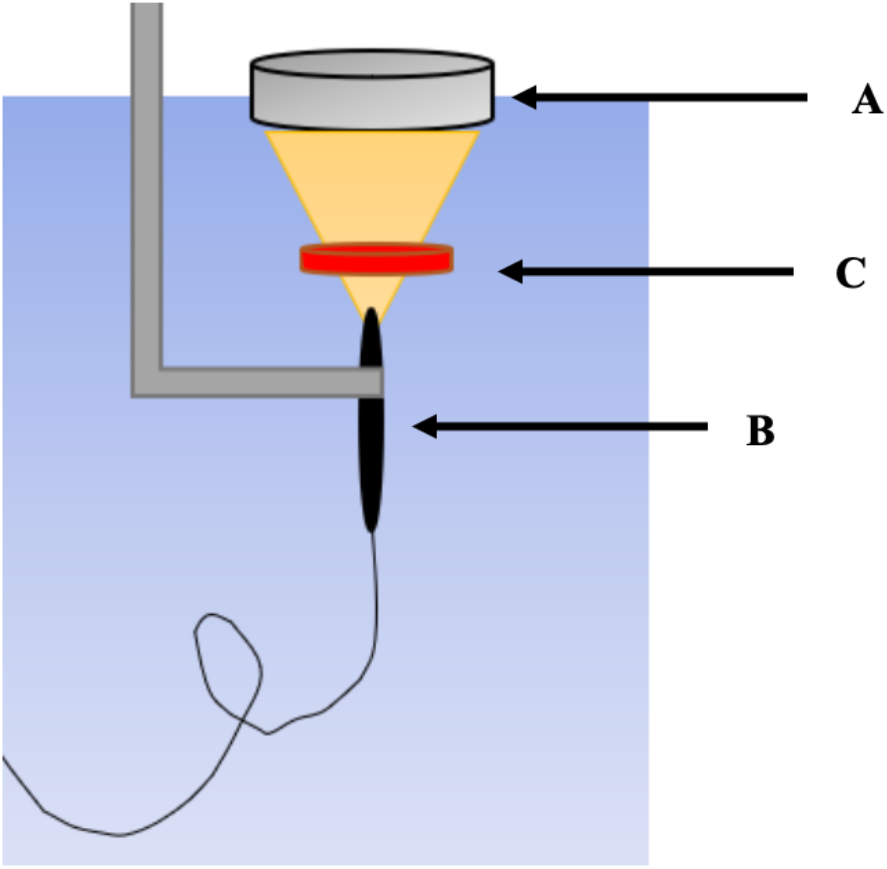
Schematic of a) transducer, b) hydrophone, and c) sample test configuration in water tank

Peak-to-peak voltage was recorded using a Tektronix TDS3012C oscilloscope. The transmit voltage was fixed and served as a reference. The measured peak-to-peak hydrophone voltage represented the received signal after passing through each pad. Attenuation in decibels was calculated as 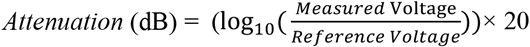

Measurements were repeated four times for each condition to confirm reproducibility, determined to be <2%.

## Results

Twenty-three sets of transmitting and non-transmitting pad pairs were created and tested. The transmitting versions were tested for their ability to allow ultrasound to pass through with minimal loss. The non-transmitting pads were tested for their ability to effectively block the ultrasound transmission, while adhering to the -40 dB loss minimum threshold, which corresponds to a factor of 10,000 in intensity. After completing the transmission tests, the transmitting version of the pads averaged -0.41 dB loss with a standard deviation of ± 0.53, which fulfills the original design requirement of less than 5% overall acoustic loss through the transmitting version of the pad. The non-transmitting version of the pad averaged -48.61 dB loss with a standard deviation of ± 4.33, which meets the original design requirement of non-transmitting pads needing to exhibit at least -40 dB loss.

Because the Dragon Skin 10 NV base material has a sound speed that is less than water, it has the potential to shift the nominal focal depth of the transducer due to refraction. Circularly symmetric ultrasound transducers,such as those used for neuromodulation, produce a focal zone with a relatively long focal extent. For example typical 62mm diameter, 650kHz transducers produce elliptically shaped focal regions with focal extents (-6dB) of between 25 and 40mm and beam widths on the order of 4mm, depending on the nominal focal depth (Schafer, et al. 2021). Separate tests were conducted to determine the effects of the pads on a transducer with an 80mm nominal focal depth. For typical transmitting pads less than 5mm thick, the shift in the axial position of the focus was less than 2mm, with no measurable change in the focal width.

In addition to meeting the design requirements for transmission, the final pad design met the visual and mechanical design requirements as well. Both types of pads are visually identical and can be casually manipulated without the user being able to distinguish between them. Figure 6 below shows an image of both types of pads, one transmitting and one non-transmitting, next to one another.

**Figure 6:**
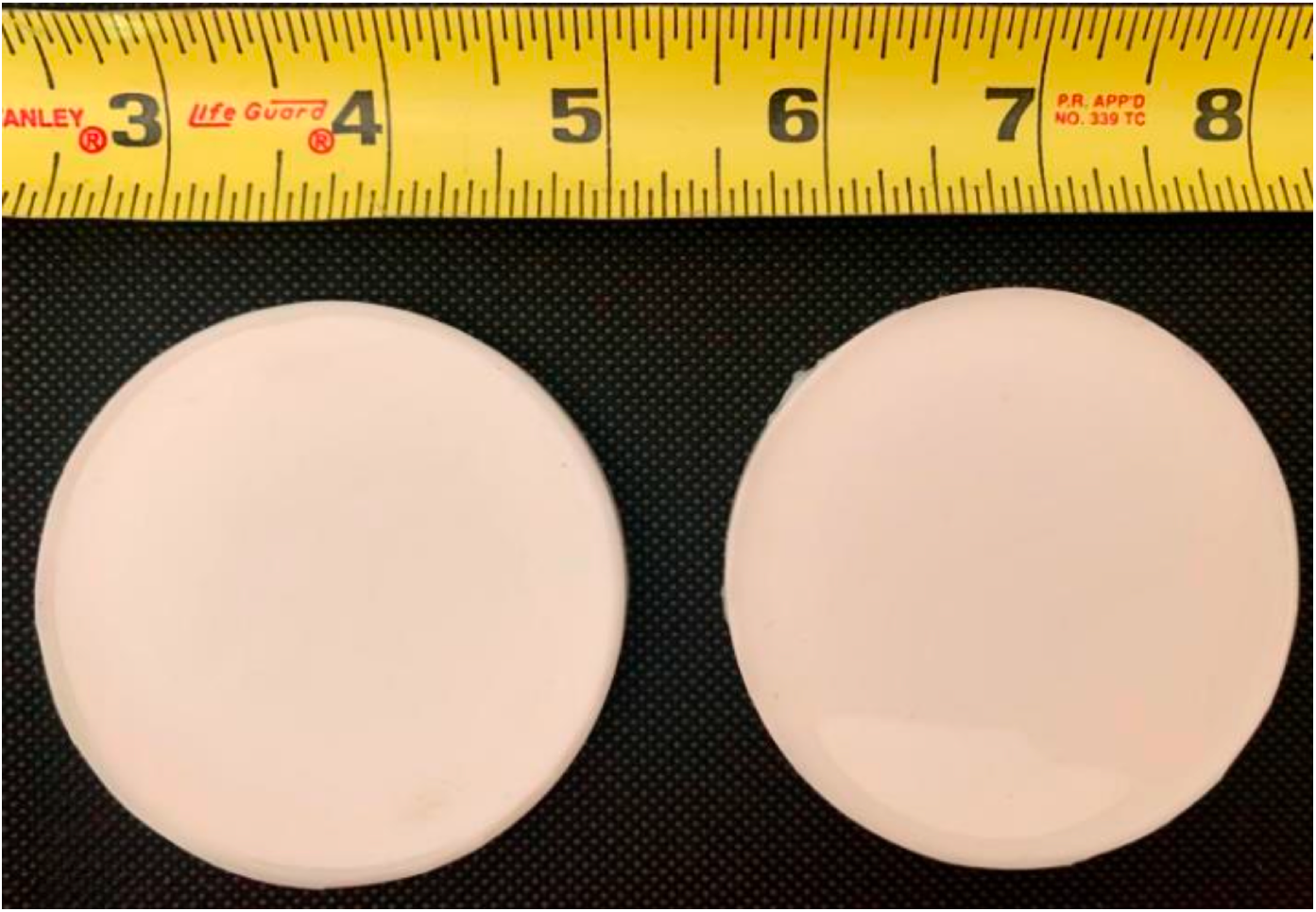
Side by side image of both transmitting and non-transmitting pads

To test whether the pads were indeed indistinguishable from one another, a set was sent to be handled under an experimental environment by an knowledgeable operator, who could not distinguish them either by physical manipulation or by visual inspection. Additionally, a participant could not distinguish between the two exposure conditions when ultrasound was applied with both pads, one after the other. Furthermore, these pads were found to be suitable for sustained reuse, as they could be sanitized with isopropyl alcohol between uses and the silicone formulation maintained its integrity under repeated cleaning.

## Discussion

The purpose of this research was to find a method of effectively blinding low-intensity (non-ablative) focused ultrasound (LIFU) for the use in neuromodulation studies. Past and current research groups have observed placebo effects in participant groups indicating an urgent need for a reliable blinding method to substantiate the data they collect. Such reliable method would ensure statistical significance and determine the modality’s (i.e. LIFU) efficacy in the treatment of neurological and psychiatric disorders. The data from both the transmission testing and the sound speed testing provided valuable insights on how the base material and final pad assembly performed in experiments with a 650 kHz LIFU transducer. Importantly, the setup employed here ensured a sufficient attenuation of ultrasound at the focal zone of the transducer, making these pads suitable for blinding in neuromodulation studies. The transmitting pads produced little attenuation or refractive effects on the ultrasound beam.

It is important to note, however, that using an opaque, dyed silicone pad—necessary to hide internal structures and keep the study blinded—prevents operators from visually checking if air bubbles are trapped between the pad and the transducer face. As pointed out earlier (see Introduction) even small air pockets can considerably weaken ultrasound energy delivered to the treated tissue volume; careful, air-free, coupling is essential to ensure adequate contact for treatment. To help prevent this, a standardized technique for applying gel should be utilized. A deposit of the spreadable gel should be placed at the center of the transducer face, and then the pad is coupled by pressing down in the center of the pad, gradually pressing, and moving outwards. This method should allow for all potential bubbles to be pushed to the outer rim from the center.

## Conclusion

LIFU is an emerging modality of neuromodulation with the potential to assist individuals suffering from various neurological and neuropsychiatric disorders. However, human studies have demonstrated inadequacies and biases due to lack of an effective method permitting both single and double blinding. The approach presented here offers a solution for effectively enabling single- and double-blind ultrasound neuromodulation studies by modifying the coupling pad used for ultrasound transmission. The final design of the pad types meets the necessary requirements and can effectively block ultrasound transmission while preserving the auditory sensations that a participant might experience during treatment. Given the sound speed characteristics of Dragon Skin 10 NV, it was determined to be a suitable base material for standard-thickness pads and it did not distort the shape of the normal focal zone of the ultrasound beam. These pads successfully provide a blinding method for LIFU neuromodulation research and, when properly implemented in clinical settings, could help improve the statistical significance of the results of critical neurological studies.

## Funding

Funding Sources: NIH T32GM008042 (to NMS), NIH F30MH136802 (to NMS)

## Disclosures

Ms. Schafer is an inventor on neuromodulation patents. Dr. Spivak and Mr. Bishay are consultants to BrainSonix Inc. Dr. Bystritsky is the CEO and shareholder of BrainSonix. Other authors report no conflicts of interest.

